# A suture-specific oocyst wall protein COWP4 is essential for excystation and infectivity, while COWP6 links wall architecture to host interaction in *Cryptosporidium parvum*

**DOI:** 10.64898/2026.05.05.722909

**Authors:** Xiaodong Wu, Jigang Yin, Wei Qi, Peng Jiang, Ying Zhang, Di Zhang, Dongqiang Wang, Guan Zhu

## Abstract

The oocyst of *Cryptosporidium* is a highly resilient transmission stage that protects sporozoites in the environment and mediates infection through a specialized opening known as the suture. While recent studies have identified *Cryptosporidium* oocyst wall proteins (COWPs) as structural components of the oocyst wall, the functional roles of individual COWPs and the molecular basis of suture biology remain poorly understood. Here, we performed a detailed characterization of two COWP family members, COWP4 and COWP6, combining immunolocalization, ultrastructural analysis, protein interaction assays, and genetic manipulation. We show that COWP4 is strictly localized to the oocyst suture, whereas COWP6 is distributed throughout the oocyst wall inner layer and enriched at the suture. COWP6 is additionally present in sporozoites and is secreted during parasite motility and host cell invasion, where it exhibits high-affinity binding to host cells. Structural analyses indicate that both proteins are cysteine-rich and likely form disulfide-stabilized architectures consistent with roles in wall assembly. Functional analyses reveal that COWP4 is essential for proper suture formation, excystation, and parasite infectivity, establishing the suture as a genetically defined gatekeeper for parasite transmission. In contrast, COWP6 functions as a multifunctional protein linking oocyst wall architecture to host interaction. We further demonstrate that COWP4 and COWP6 interact, suggesting coordinated assembly within the oocyst wall. Together, these findings provide a functional dissection of the *Cryptosporidium* oocyst suture and reveal distinct, non-redundant roles for individual COWPs. This work advances our understanding of oocyst wall biology and identifies COWP4 as an essential determinant of parasite transmission that may be targeted to disrupt the infectious cycle.

**Author summary:** *Cryptosporidium* is a major cause of diarrheal disease worldwide and is transmitted through environmentally resistant oocysts that protect infectious stages of the parasite. A defining feature of the oocyst is a specialized seam-like structure, called the suture, through which parasites exit to initiate infection. Although oocyst wall proteins (COWPs) have been identified, how individual proteins contribute to the structure and function of the oocyst wall, particularly the suture, has remained unclear. In this study, we investigated two oocyst wall proteins, COWP4 and COWP6, and uncovered distinct roles for each. We found that COWP4 is specifically localized to the suture and is essential for its proper formation, enabling parasites to exit the oocyst and establish infection. In contrast, COWP6 is distributed more broadly in the oocyst wall and also functions beyond it, being secreted during parasite movement and interacting with host cells. These findings reveal that the oocyst wall is composed of specialized proteins with non-redundant functions and identify the suture as a critical control point for parasite transmission. Understanding these mechanisms may help guide the development of new strategies to block infection by disrupting oocyst integrity or parasite release.

**Highlights:**

- COWP4 is a suture-specific protein essential for oocyst excystation and infectivity
- Genetic disruption of COWP4 produces non-infectious oocysts with impaired viability
- The oocyst suture is a genetically defined gatekeeper for parasite transmission
- COWP6 localizes to the wall inner layer and is enriched at the suture
- COWP6 is secreted by sporozoites and binds host cells with nanomolar affinity
- · COWP4 and COWP6 directly interact, linking wall architecture to host interaction

## Introduction

*Cryptosporidium* is a genus of diarrheal protozoan parasites that infect humans and animals, contributing substantially to morbidity and mortality in vulnerable populations, including young children, the elderly, and immunocompromised individuals [1,2]. Human infections are primarily caused by the zoonotic species *C. parvum* and the anthroponotic species *C. homini*s, which are also major contributors to recurrent waterborne outbreaks worldwide [3,4]. Transmission occurs via the fecal–oral route through environmentally resistant oocysts, which contain four sporozoites enclosed within a robust wall [2]. These subspherical oocysts, approximately 5 μm in diameter, are difficult to remove by conventional sedimentation and filtration during water treatment. In addition, the oocyst wall is highly resistant to commonly used disinfectants, allowing persistence in water and food matrices and posing a significant public health risk [5–7]. A detailed understanding of the structural and molecular features of the oocyst wall is therefore both biologically important and relevant for developing strategies to interrupt transmission.

Earlier ultrastructural, biochemical, and immunolabeling studies have established that the *Cryptosporidium* oocyst wall is a multilayered structure with distinct molecular compositions [8–11]. Integrating findings from these studies, the wall can be described as comprising four major layers: (i) a moderately electron-dense outer veil (OV), also referred to as a glycocalyx layer of approximately 8.5 nm; (ii) an electron-translucent lipid hydrocarbon layer (LHL; ∼4.0 nm); (iii) an electron-dense oocyst wall central layer (OWCL; ∼13 nm); and (iv) a moderately electron-dense oocyst wall inner layer (OWIL; 25–40 nm) (Fig. 1A, inset). The OV consists mainly of glycoproteins and glycolipids and can be readily removed by chlorine treatment or cesium chloride centrifugation. The OWCL is resistant to detergents and proteases, although surface-exposed proteins can be removed by proteolysis. After sonication and protease digestion, the OWCL exhibits a bilayer architecture. In contrast, the OWIL is proteinaceous, enriched in glycoproteins, and susceptible to protease digestion. The OWIL is also associated with fibrous glycoproteins, some of which tether the enclosed sporozoites. The OWCL is resistant to detergent and proteases, although proteases can remove its surface-exposed proteins. With sonication and protease-digestion of excysted oocyst walls, the rigid OWCL displays a bilayer architecture. The OWIL is proteinaceous, rich in glycoproteins and susceptible to protease digestion. OWIL is also attached by fibrous glycoproteins, some of which tether the sporozoites [9]. Although the oocyst surface appears smooth and continuous, an electron-translucent structure known as the suture is embedded within the inner layers and extends along approximately one-third of the oocyst circumference at one pole [8,11]. The suture functions as a gate through which sporozoites are released during excystation following ingestion by a host.

**Fig 1.**
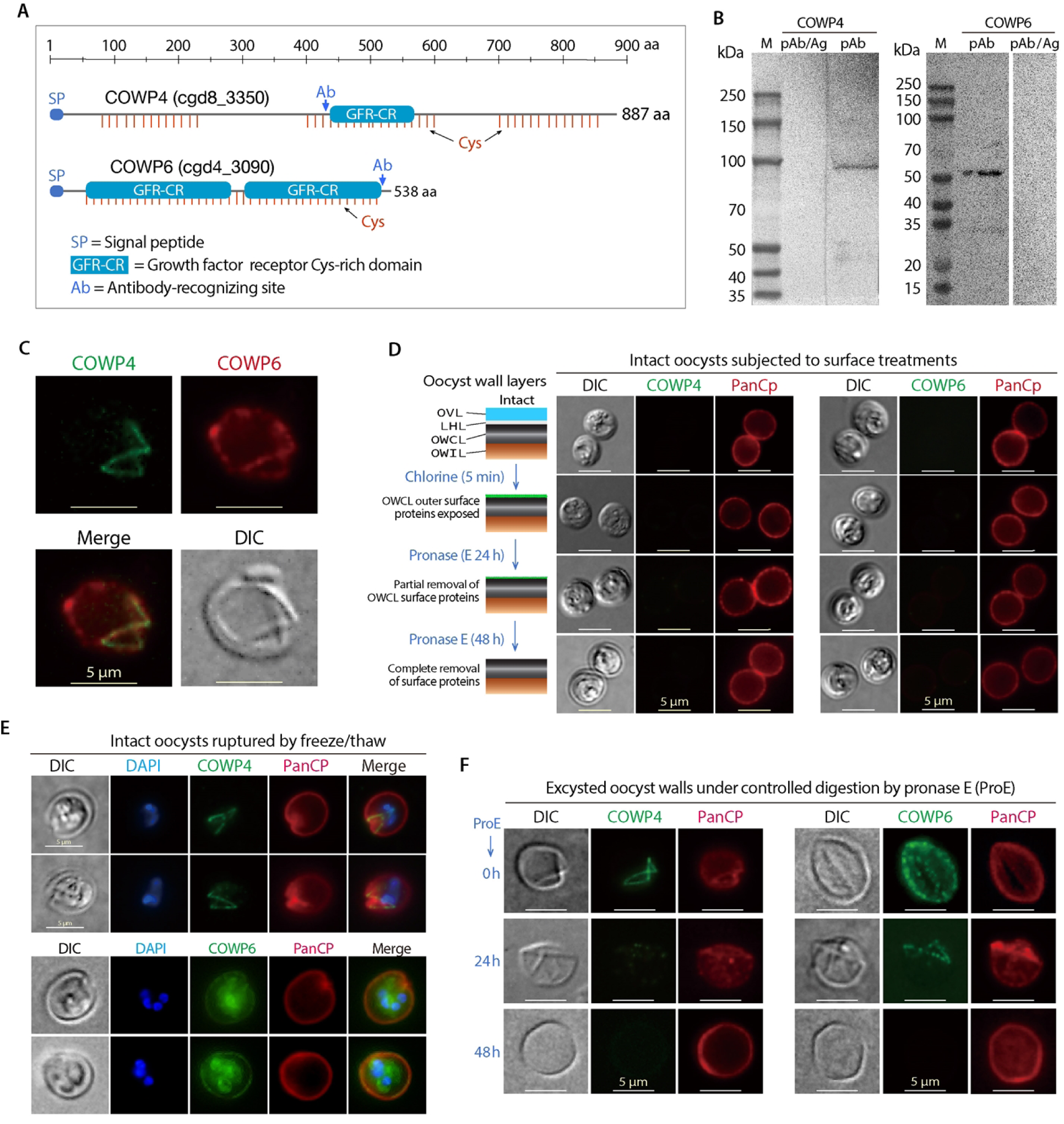
Differential localization of *C. parvum* oocyst wall proteins COWP4 and COWP6 at the suture and within the oocyst wall inner layer. **(A)** Domain organization of COWP4 and COWP6. Both proteins contain an N-terminal signal peptide (SP) and one or more growth factor receptor cysteine-rich (GFR-CR) domains. Cysteine residues are indicated by red vertical lines. Peptide regions used for antibody (Ab) production are marked. See also S1 Fig for AlphaFold-predicted structural models. **(B)** Western blot analysis of COWP4 and COWP6 in *C. parvum* oocyst extracts using affinity-purified antibodies. Each antibody detected a single band at the expected molecular weight (lane labeled pAb), and signal detection was abolished by pre-incubation with the corresponding antigen (lane labeled pAb/Ag), confirming specificity. **(C)** Dual-label immunofluorescence assay (IFA) showing localization of COWP4 at the oocyst suture and COWP6 across the oocyst wall in excysted oocyst wall preparations. **(D)** IFA of intact oocysts subjected to surface treatments. Neither COWP4 nor COWP6 was detectable in intact oocysts with preserved outer veil (OV), or in oocysts treated with chlorine and pronase E (0.7 U/mL at 25 °C) to remove the OV and expose proteins on the outer surface of the oocyst wall central layer (OWCL). **(E)** IFA of oocysts ruptured by freeze–thaw cycles. COWP4 localizes to the open suture, whereas COWP6 is detected across the oocyst wall and within enclosed sporozoites. **(F)** Controlled proteolytic digestion of excysted oocyst walls with pronase E (0.7 U/mL at 25 °C). In untreated samples (0 h), COWP4 shows suture-specific labeling, while COWP6 is distributed across the oocyst wall. After 24 h digestion, COWP4 signals at the suture are largely eliminated, whereas COWP6 signals become apparent at the suture but are reduced in the wall. After 48 h digestion, signals for both proteins are completely abolished. These results indicate that COWP4 is suture-specific, whereas COWP6 is present in the suture, the oocyst wall inner layer (OWIL), and sporozoites, and exhibits greater resistance to proteolytic removal at the suture. In panels D–F, samples were co-labeled with a pan-C. parvum (PanCp) antibody. In panel E, nuclei were counterstained with DAPI.

Despite its importance, the molecular composition and organization of the oocyst wall remain incompletely understood. *Cryptosporidium* oocyst wall protein 1 (COWP1; cgd6_2090) was the first experimentally characterized component of the inner wall layer [12]. COWP1 is characterized by a periodic distribution of cysteine residues arranged in tandem arrays of type I and type II domains. Subsequent genome mining identified eight additional COWP family members (COWP2 to COWP9) containing similar cysteine-rich motifs [13]. These proteins are thought to contribute to the structural integrity of the oocyst wall, potentially through extensive disulfide bond formation. Additional studies have identified glycoproteins such as CpGP15 in the outer veil, along with other glycoproteins distributed in inner layers [9]. More recently, transgenic *C. parvum* parasites expressing fluorescently tagged COWPs revealed distinct subcellular localizations, with some proteins enriched at the suture and others distributed throughout the wall [14]. Notably, genetic disruption of COWP8 showed that it is dispensable for oocyst formation and transmission, suggesting functional diversification within the COWP family [14].

In the present study, we performed a detailed characterization of COWP4 and COWP6 to define their roles in oocyst wall organization and function. We show that COWP4 is specifically localized to the oocyst suture, whereas COWP6 is a multifunctional protein present in both the suture and the inner wall layer. In addition, COWP6 is detected in the residual body and sporozoites, is secreted during parasite motility and invasion, and binds to host cells with high affinity. Importantly, gene disruption demonstrates that COWP4 is essential for suture formation, excystation, and parasite infectivity, establishing a critical role for this protein in parasite transmission. These findings provide new insights into the functional specialization of COWPs and the molecular basis of oocyst wall architecture.

## RESULTS

### COWP4 is a suture-specific oocyst wall protein, whereas COWP6 is distributed in both the suture and the oocyst wall inner layer

The COWP family is characterized by a high content of cysteine residues arranged in tandem arrays, typically spaced at intervals of approximately 10–12 amino acids [13]. COWP4 (cgd8_3350; 887 aa) contains 44 cysteine residues (4.96%) organized into three distinct clusters and includes a central growth factor receptor cysteine-rich (GFR-CR) domain (Fig. 1A). Similarly, COWP6 (cgd4_3090; 538 aa) is highly cysteine-rich (7.43%; 40 residues) and contains two GFR-CR domains. Both proteins possess N-terminal signal peptides, consistent with entry into the secretory pathway. The periodic distribution of cysteine residues and the presence of GFR-CR domains suggest that COWP4 and COWP6 adopt disulfide-stabilized architectures capable of forming crosslinked structural frameworks within the oocyst wall.

AlphaFold-predicted structural models indicate that both proteins adopt elongated, modular conformations composed of β-strand-rich elements connected by disulfide bonds (S1 Fig). Compared to COWP6, COWP4 displays a more complex architecture, with flexible loop regions separating the cysteine-rich clusters. These features are consistent with roles in protein–protein interactions and in the assembly and mechanical stability of the oocyst wall.

To determine the sublayer localization of COWP4 and COWP6, we generated affinity-purified polyclonal antibodies against synthetic peptide epitopes (Fig. 1A). Each antibody detected a single band in oocyst extracts at the expected molecular weight, and the signals were abolished by peptide competition, confirming specificity (Fig. 1B).

Immunofluorescence analysis of excysted oocyst walls revealed that COWP4 is strictly localized to the suture, whereas COWP6 is distributed throughout the oocyst wall and enriched at the suture (Fig. 1C). Neither protein was detectable on the surface of intact oocysts, including those treated with chlorine or protease to remove the outer veil and expose the outer surface of the central layer (Fig. 1D). However, following mechanical disruption of the oocyst wall by freeze–thaw treatment, COWP4 labeling remained restricted to the suture, whereas COWP6 signals were observed along the entire wall and within enclosed sporozoites (Fig. 1E).

To further probe the relative organization of COWP4 and COWP6 at the suture, we performed controlled proteolytic digestion of excysted oocyst walls using low concentrations of pronase E under mild conditions (Fig. 1F). Prior to digestion (0 h), the distributions of COWP4 and COWP6 were consistent with those described above. After prolonged digestion (48 h), signals for both proteins were completely lost, indicating full removal. Notably, at an intermediate time point (24 h), COWP4 signals were no longer detectable, whereas COWP6 signals were selectively retained at the suture but largely absent from the oocyst wall inner layer.

This differential sensitivity to proteolysis suggests that COWP4 is more accessible to enzymatic digestion, whereas a subset of COWP6 molecules, particularly those associated with the suture, are more resistant. These observations are consistent with a model in which COWP4 and COWP6 are differentially organized at the suture, with COWP4 occupying a more exposed position and COWP6 contributing to a less accessible, potentially underlying structural layer. While this interpretation does not resolve the precise molecular arrangement, it supports the notion of a stratified or hierarchically organized protein architecture within the suture region. The selective retention of COWP6 at the suture, but not in the inner wall, further suggests that COWP6 adopts distinct structural contexts in these two locations.

### Immunogold labeling confirms preferential localization of COWP4 to the suture

To quantitatively assess the spatial distribution of COWP4 and COWP6 on the oocyst wall, we performed immunogold labeling on excysted oocyst walls followed by transmission electron microscopy (Fig. 2A–C; S2 Fig). Consistent with the immunofluorescence results, both proteins were highly enriched at the open suture (Fig. 2A,B). COWP4 showed a markedly higher proportion of gold particles at the suture, accounting for 85.2% of the total particles detected on the oocyst wall (Fig. 2C). COWP6 also displayed preferential localization to the suture, accounting for 61.8% of the total gold particles.

**Fig. 2.**
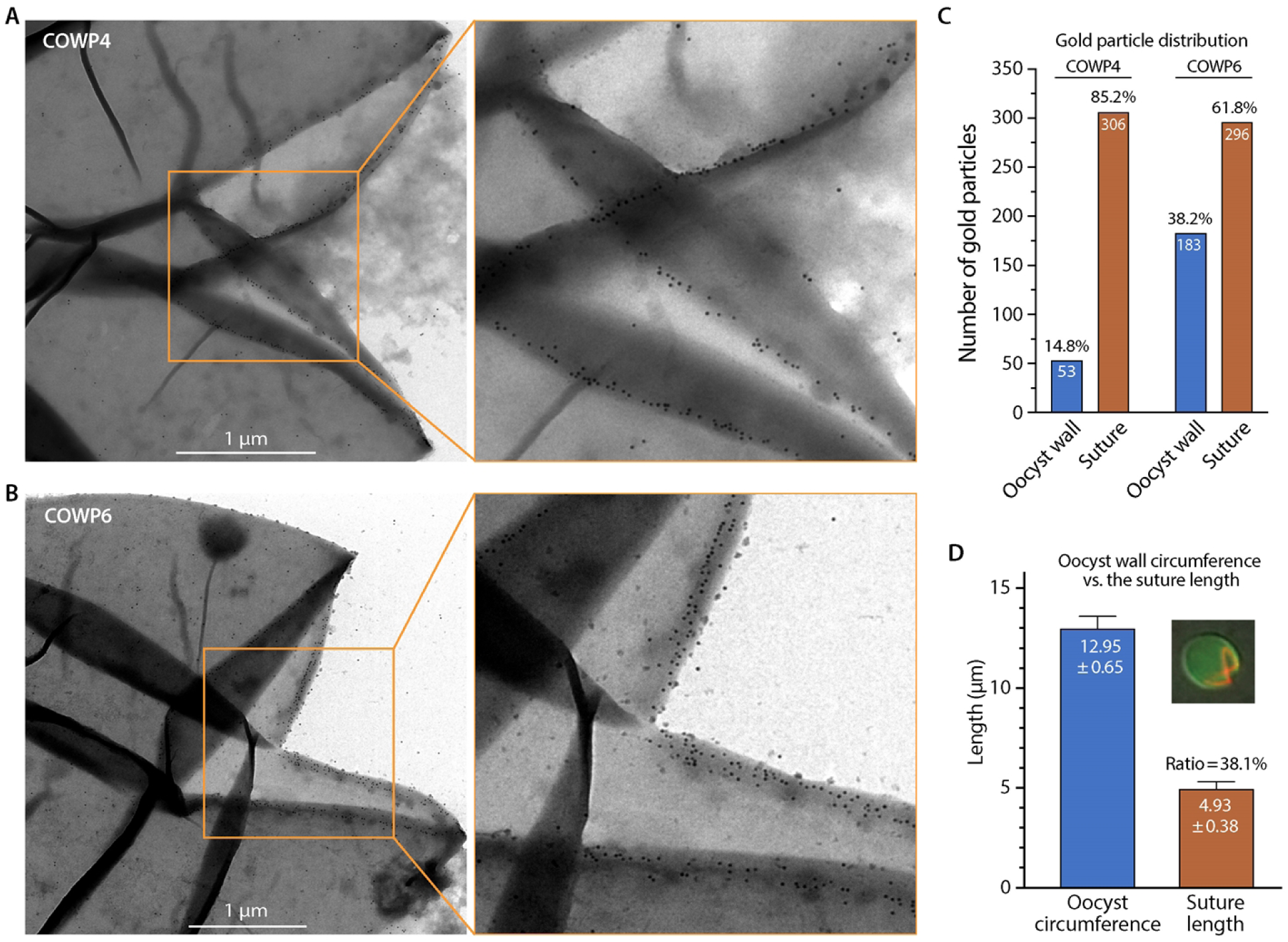
Immunogold electron microscopy (IEM) analysis of COWP4 and COWP6 in the *C. parvum* oocyst wall. (A–B) Representative IEM micrographs showing the distribution of colloidal gold-labeled COWP4 **(A)** and COWP6 **(B)** on excysted oocyst walls. COWP4 labeling is concentrated at the suture edges, whereas COWP6 labeling is detected both at the suture and along other regions of the oocyst wall. Enlarged views of boxed regions are shown to highlight labeling patterns. See also **S2 Fig** for additional IEM images. **(C)** Quantification of gold particles corresponding to COWP4 and COWP6 at the suture and other regions of the oocyst wall. Values are presented as the number and proportion of particles detected in each region. **(D)** Measurements of oocyst wall geometry based on IFA-labeled oocyst walls, showing the average suture length (4.93 ± 0.38 μm) relative to the total oocyst wall circumference (12.95 ± 0.65 μm), corresponding to 38.1% of the perimeter.

Given that COWP6 is also distributed in the oocyst wall inner layer, its relatively high proportion of labeling at the suture was somewhat unexpected. This apparent enrichment may, in part, reflect limited accessibility of antibodies and gold particles to the inner layer in flattened oocyst wall preparations used for immunogold labeling, which could bias detection toward more exposed regions such as the suture.

Measurements of oocyst wall geometry showed that the suture has an average length of 4.93 ± 0.38 μm, corresponding to approximately 38.1% of the total oocyst wall circumference (12.95 ± 0.65 μm) (Fig. 2D). Despite occupying a minority of the oocyst perimeter, the suture contains a disproportionately large fraction of the immunogold labeling for both proteins, particularly for COWP4. Although the exact surface area of the suture cannot be determined, this disparity strongly supports a preferential accumulation of COWP4, and to a lesser extent COWP6, at the suture.

Together, these results confirm that COWP4 is a suture-specific protein, while COWP6 is present both at the suture and across the oocyst wall inner layer.

### Differential expression of COWP4 and COWP6 during parasite development and oocyst wall formation

To examine the temporal expression and localization of COWP4 and COWP6 during parasite development, we performed immunofluorescence analysis using specific antibodies against each protein (green), together with a pan-C. parvum antibody (PanCp; red) to visualize parasite cells (Fig. 3).

**Fig 3.**
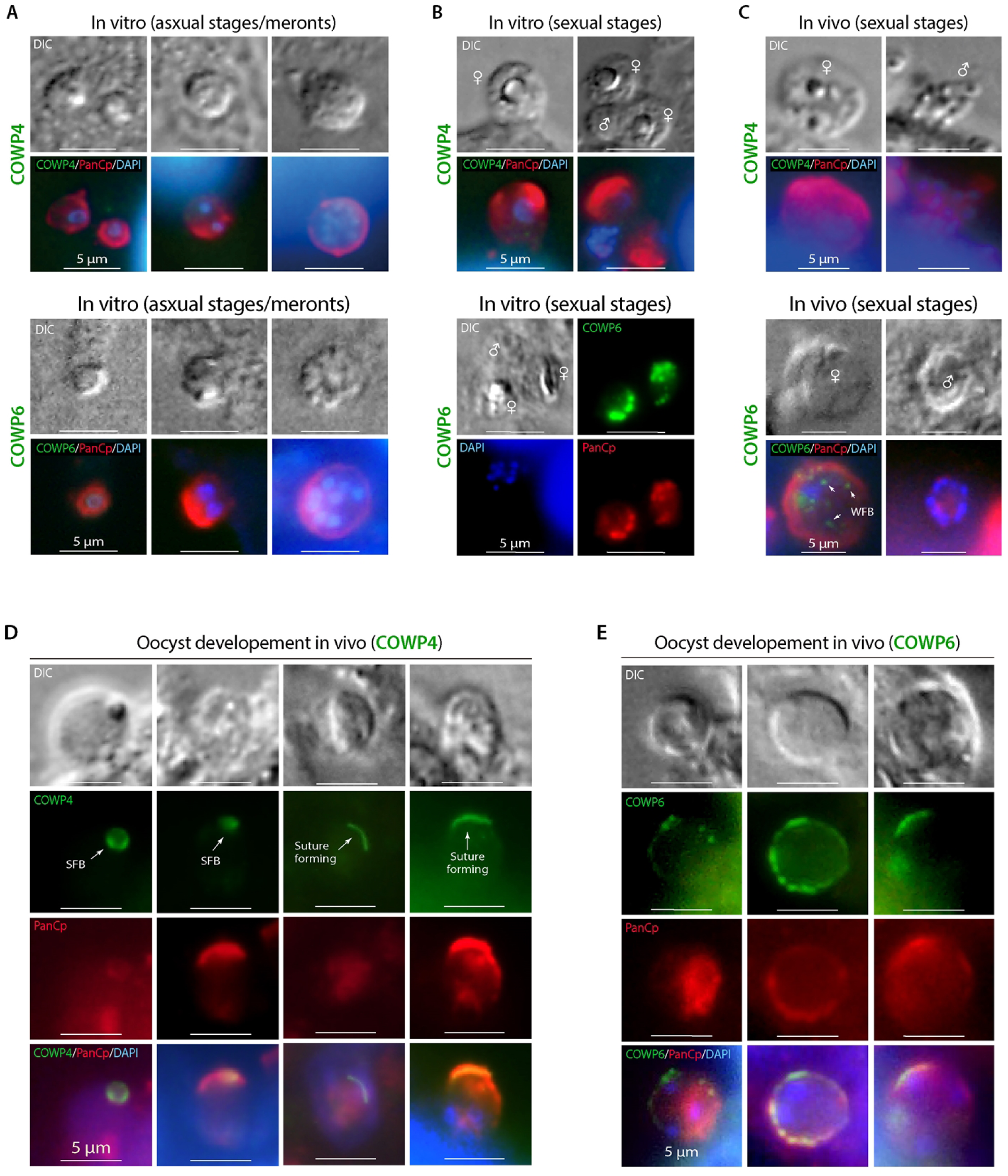
Immunolabeling of COWP4 and COWP6 during intracellular development of *C. parvum*, including asexual and sexual stages and oocyst formation. Samples were prepared in vitro (A–B) using infected HCT-8 cells or in vivo (C–E) from infected IFN-γ-KO mice. Parasites were labeled with antibodies against COWP4 or COWP6 (green) and co-stained with a pan-*C. parvum* (PanCp) antibody (red) and DAPI for nuclei (blue). **(A)** In asexual stages (meronts) obtained in vitro, both COWP4 and COWP6 were undetectable. **(B)** In sexual stages obtained in vitro, both proteins were absent in male cells. COWP4 was also absent in female cells (upper panel), whereas COWP6 was detected as multiple punctate granules in female cells (lower panel), consistent with wall-forming bodies (WFBs). **(C)** In sexual stages obtained in vivo, both COWP4 and COWP6 were absent in male cells. COWP4 was also absent in female cells (upper panel), whereas COWP6 showed scattered granular labeling in female cells (lower panel), consistent with WFB-like structures. **(D)** During oocyst formation in vivo, COWP4 was detected as either single round structures or curved line-like signals. The round structures are tentatively interpreted as suture-forming bodies (SFBs), representing a vesicular compartment distinct from WFBs, while the curved signals correspond to the developing suture. **(E)** During oocyst formation in vivo, COWP6 exhibited a granular distribution surrounding parasite cells, consistent with its deposition into the oocyst wall inner layer via WFBs.

Under in vitro culture conditions, *C. parvum* undergoes asexual replication (merogony) and sexual differentiation into macrogamonts (female) and microgamonts (male), but does not progress to fertilization and oocyst formation [15,16]. In these samples, both COWP4 and COWP6 were undetectable in asexual stages, showing no or near-background signals in meronts (Fig. 3A). COWP4 was also not detected in male and female sexual stages (Fig. 3B, upper panel), indicating that it is not expressed during asexual or pre-fertilization sexual development.

In contrast, COWP6 was absent in male cells but showed strong punctate labeling in female cells (Fig. 3B, lower panel). These signals appeared as multiple discrete granules, consistent with the morphology of previously described wall-forming bodies (WFBs), which mediate the transport of oocyst wall components. These observations indicate that COWP6 is expressed in female cells prior to fertilization and is likely trafficked via WFBs during early stages of oocyst wall assembly.

To investigate later developmental stages, we analyzed intestinal samples from infected IFN-γ-KO mice, in which the parasite completes its life cycle in vivo. In these samples, COWP4 remained undetectable in male cells and in some female cells that likely represent pre-fertilization macrogamonts (Fig. 3C, upper panel). Similarly, COWP6 was absent in male cells but showed scattered granular labeling in female cells (Fig. 3C, lower panel) (Fig. 3C, lower panel).

During oocyst formation in vivo, COWP4 was detected in two distinct forms: as single round structures and as curved line-like signals (Fig. 3D). The curved structures are consistent with the morphology of the developing suture. Notably, the round COWP4-positive structures differed from the multi-granular WFBs observed for COWP6. We therefore tentatively interpret these structures as a distinct vesicular compartment involved in suture assembly and refer to them here as suture-forming bodies (SFBs). COWP4 labeling was restricted to these intermediate stages and was not detected in mature oocysts, consistent with its localization to the suture, which becomes inaccessible following completion of the oocyst wall.

In parallel, COWP6 labeling in vivo was observed as multiple granules surrounding developing parasite cells (Fig. 3E), consistent with ongoing oocyst wall formation. These granules likely correspond to WFBs delivering COWP6 to the forming inner wall layer. Similar to COWP4, COWP6 was not detectable in mature oocysts, likely due to limited accessibility of antibodies to the inner wall layer after wall completion

Together, these results indicate that COWP6 is expressed in female cells prior to fertilization and trafficked via WFBs to contribute to oocyst wall formation, whereas COWP4 is expressed after fertilization and is specifically associated with suture formation, potentially via a distinct vesicular pathway. These findings reveal differential timing and trafficking mechanisms for COWP family proteins during oocyst wall assembly.

### COWP4 is dispensable for intracellular development but essential for excystation and infectivity

To investigate the biological roles of COWP4 and COWP6, we attempted gene disruption using a CRISPR/Cas9-based strategy to insert an mNeonGreen–Nluc–Neo cassette into the respective loci (Fig. 4A). Despite multiple attempts, we were unable to obtain COWP6-null parasites, suggesting that COWP6 may be essential for parasite viability.

**Fig. 4.**
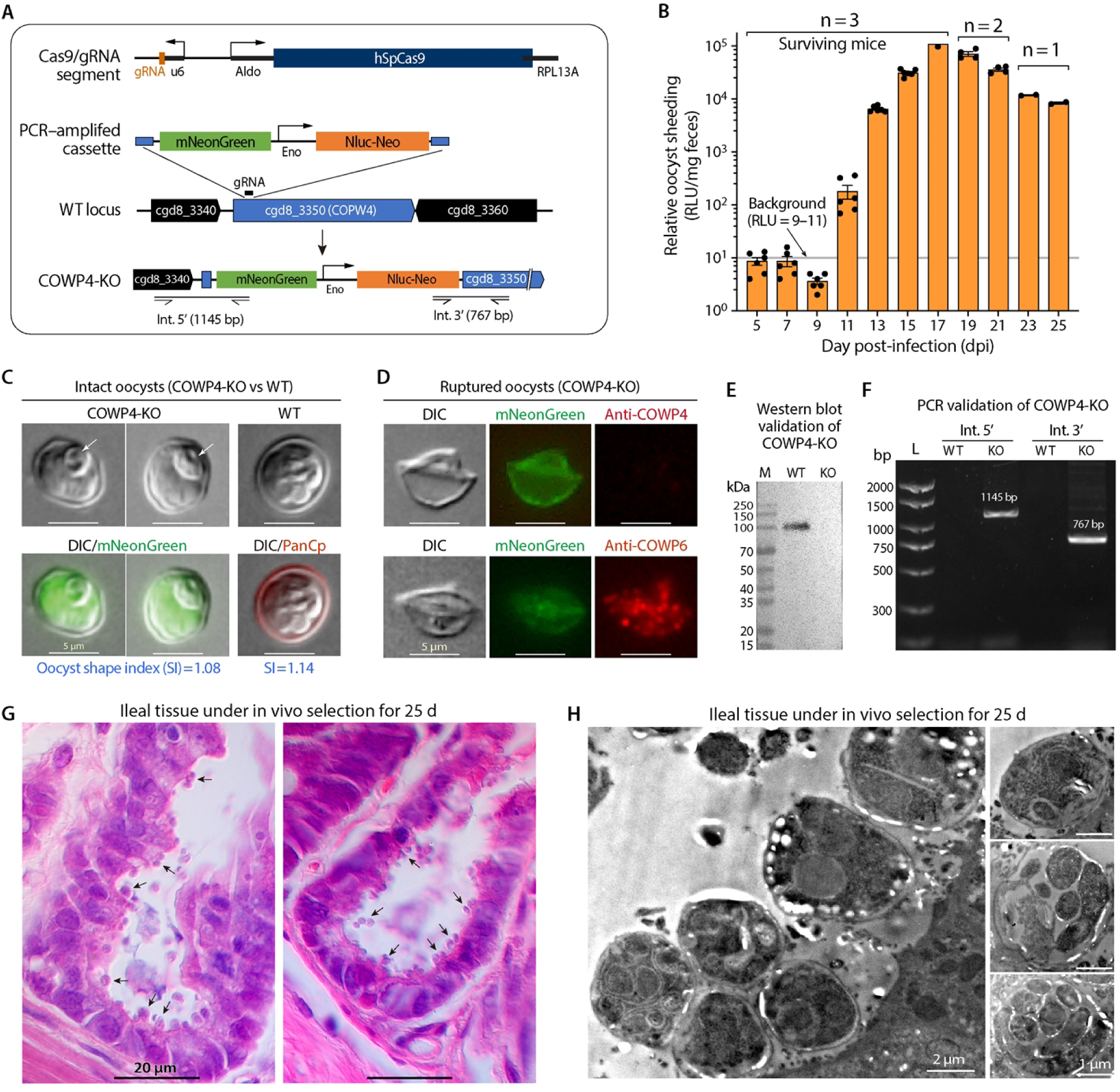
Generation and validation of the C. parvum COWP4 knockout (COWP4-KO) parasite. **(A)** Schematic of CRISPR/Cas9-mediated disruption of the COWP4 locus by insertion of the mNeonGreen–Nluc–Neo cassette. Positions of gRNA and PCR primers used for validation are indicated. **(B)** In vivo selection of COWP4-KO parasites in IFN-γ-KO mice (n = 3), monitored by luciferase activity in fecal samples. A prolonged prepatent period was observed, with detectable oocyst shedding beginning at 11 dpi. **(C)** Representative images of COWP4-KO oocysts (green fluorescence) compared to WT oocysts labeled with PanCp antibody. COWP4-KO oocysts are morphologically similar but slightly more elongated. Arrows indicate a ring-like internal structure observed in COWP4-KO oocysts. See also S3 and S4 Figs. **(D)** Immunofluorescence validation of COWP4 disruption. Ruptured COWP4-KO oocysts are not labeled by anti-COWP4 antibody but remain positive for anti-COWP6 labeling. **(E)** Western blot analysis confirming absence of COWP4 protein in COWP4-KO oocysts. **(F)** PCR validation of cassette integration at the COWP4 locus using primers flanking the 5′ and 3′ junctions. **(G)** Histopathology (H&E staining) showing parasite presence in ileal epithelia during in vivo selection. **(H)** Transmission electron microscopy showing intracellular developmental stages in infected ileal tissue.

In contrast, disruption of the COWP4 locus was successful (Fig. 4B–H). During in vivo selection in three IFN-γ-KO mice, a prolonged prepatent period of approximately 10 days was observed, with luciferase-positive oocyst shedding detected from 11 days post-infection (dpi) and persisting for over two weeks (Fig. 4B). This is in contrast to wild-type (WT) parasites, which typically begin shedding oocysts at 3–5 dpi [17,18]. Despite the delayed onset, the transgenic parasites retained virulence, resulting in mortality in two mice at 18 and 22 dpi, respectively (Fig. 4B). Approximately 2.5 × 10^8^ oocysts were collected during this period (F1 generation), and nearly all exhibited mNeonGreen fluorescence (Fig. 4C; S3 Fig), indicating efficient integration of the cassette.

Disruption of COWP4 expression was confirmed by immunofluorescence and western blot analysis. Ruptured oocyst walls from COWP4-KO parasites were not labeled by anti-COWP4 antibody but remained positive for anti-COWP6 labeling (Fig. 4D), demonstrating specific loss of COWP4. Western blot analysis detected COWP4 in WT oocyst lysates but not in COWP4-KO samples (Fig. 4E). Correct integration of the mNeonGreen–Nluc–Neo cassette was further verified by PCR amplification across the 5′ and 3′ junctions, yielding products of the expected sizes (1,145 bp and 767 bp) in COWP4-KO parasites but not in WT controls (Fig. 4F).

Morphologically, COWP4-KO oocysts appeared broadly similar to WT oocysts, although they were slightly more elongated (5.30 × 4.68 μm vs. 5.25 × 4.88 μm for WT), corresponding to shape indices of 1.13 and 1.08, respectively (Fig. 4C; S4 Fig). Scanning electron microscopy revealed no obvious differences in surface texture between WT and COWP4-KO oocysts (S5 Fig). Notably, COWP4-KO oocysts frequently contained a distinct ring-like internal structure that was not observed in WT oocysts (Fig. 4C, arrows). Given that COWP4 is associated with a single vesicular structure during development, this feature may represent a residual structure arising from defective suture assembly in the absence of COWP4, although its precise origin remains to be determined.

Histopathological analysis and transmission electron microscopy revealed heavy parasite burdens in the ileal epithelia of the surviving mouse at 25 dpi (Fig. 4G,H), indicating that COWP4-null parasites are capable of establishing prolonged infection and completing intracellular development in immunodeficient hosts.

Despite successful production of COWP4-KO oocysts, these parasites exhibited profound defects in transmission-related processes. COWP4-KO oocysts failed to undergo excystation in vitro (0% excystation after 1–2 h incubation, compared to 88.2% for WT; Fig. 5A), demonstrating that COWP4 is essential for sporozoite release. In addition, COWP4-KO oocysts showed markedly reduced viability during storage at 4 °C. Relative viability declined to 78.5%, 33.2%, 17.3%, and 0% after 7, 30, 60, and 90 days, respectively, as determined by qRT-PCR (Fig. 5B), and complete loss of viability at 90 days was further confirmed by ATP assay (Fig. 5C). These findings suggest that deletion of COWP4 compromises pathways required for long-term oocyst survival.

**Fig. 5.**
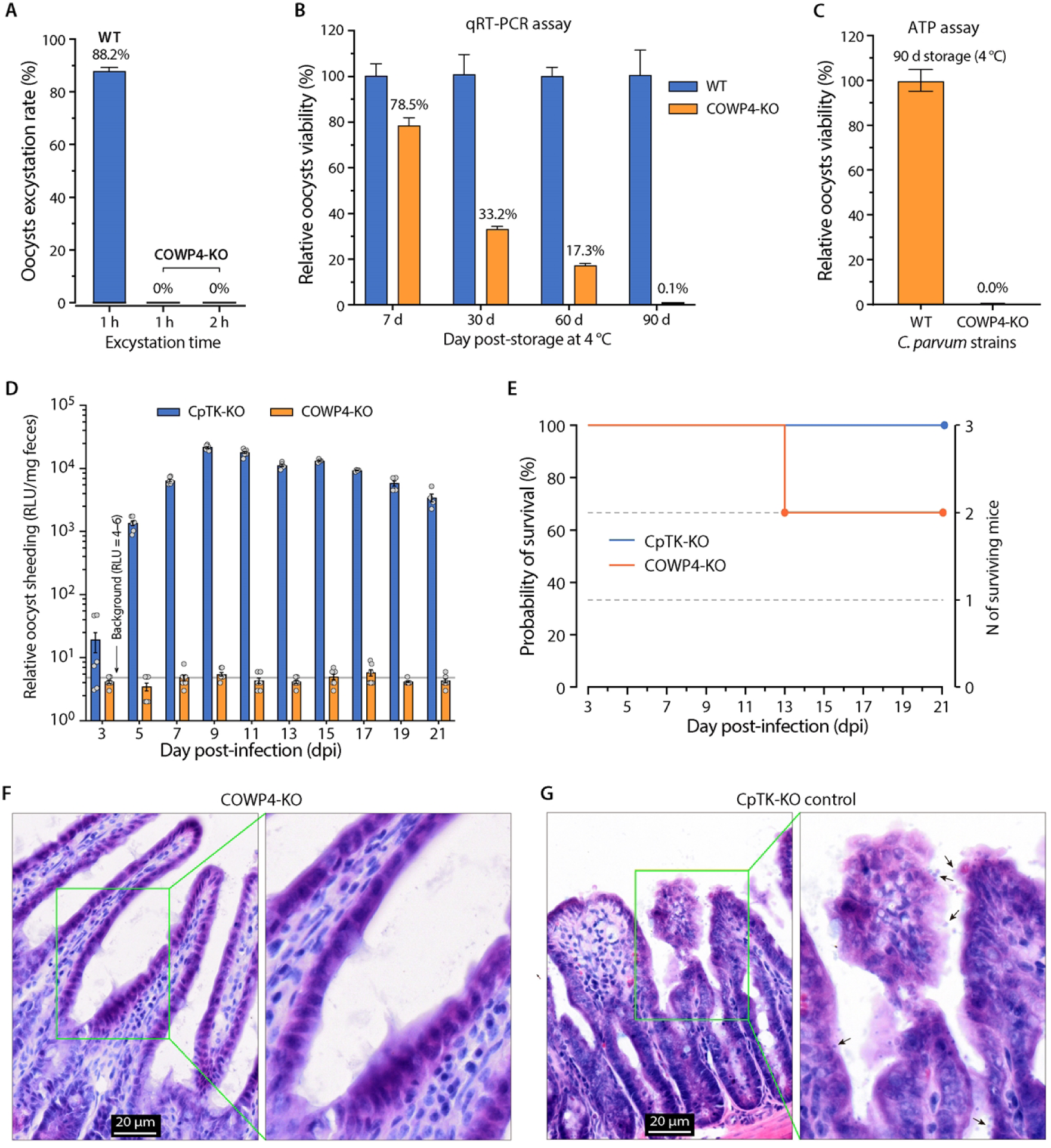
Phenotypic characterization of COWP4-KO oocysts. **(A)** COWP4-KO oocysts fail to undergo in vitro excystation, showing 0% excystation compared to 88.2% for WT oocysts. **(B)** Decline in viability of COWP4-KO oocysts during storage at 4 °C, measured by qRT-PCR detection of 18S rRNA. **(C)** ATP-based assay confirming complete loss of viability of COWP4-KO oocysts after 90 days of storage. **(D)** COWP4-KO oocysts fail to establish infection in IFN-γ-KO mice, as indicated by absence of luciferase signals and oocyst shedding. CpTK-KO parasites serve as a positive control. **(E)** Survival curves of infected mice, showing no mortality in the COWP4-KO group. **(F–G)** Histopathology (H&E staining) of ileal tissue showing absence of parasites in COWP4-KO-infected mice (F) and heavy infection in CpTK-KO controls (G).

Consistent with these defects, COWP4-KO oocysts failed to establish infection in IFN-γ-KO mice, as indicated by the absence of luciferase signals and oocyst shedding (Fig. 5D). In contrast, control CpTK-KO parasites readily established infection and shed oocysts from 3 dpi onward. Infection with COWP4-KO oocysts resulted in no mortality, whereas CpTK-KO infection caused mortality in one-third of animals (Fig. 5E). Histopathological analysis confirmed the absence of parasites in mice inoculated with COWP4-KO oocysts, while heavy infection was observed in CpTK-KO controls (Fig. 5F,G).

Collectively, these results demonstrate that COWP4 is dispensable for intracellular development and oocyst formation, but is essential for excystation, infectivity, and environmental persistence of oocysts.

### COWP6 is a multifunctional oocyst wall protein that is also secreted and binds host cells

Immunofluorescence analysis showed that COWP6 is present not only in the oocyst wall but also in sporozoites and residual bodies (RBs) (Fig. 6A, left panel). In contrast, COWP4 was largely restricted to the suture, with only weak signals occasionally observed in sporozoites and no detectable signal in RBs (Fig. 6A, right panel).

**Fig. 6.**
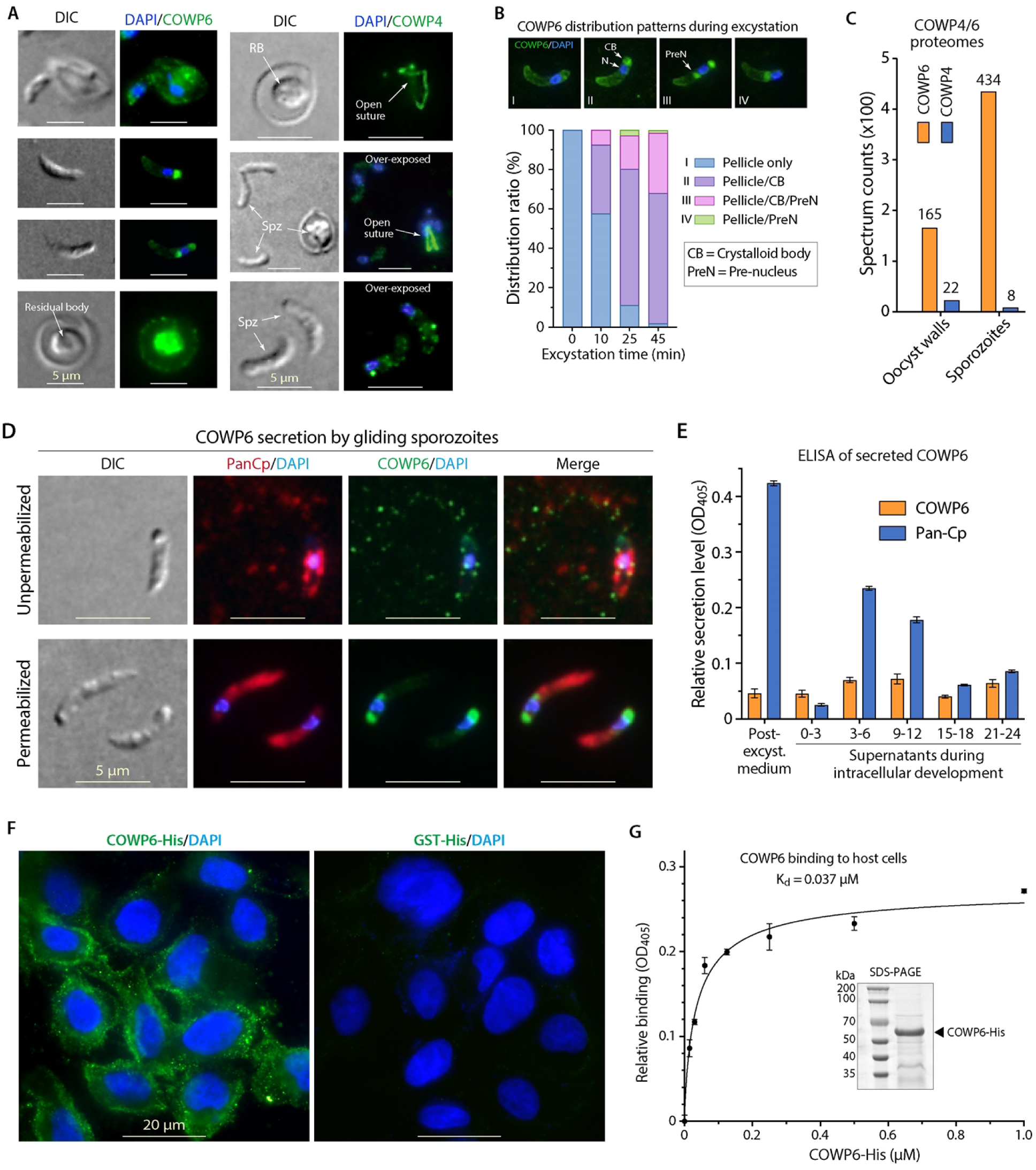
COWP6 is a multifunctional protein present in sporozoites, secreted during motility and development, and capable of binding host cells. **(A)** Immunofluorescence analysis showing COWP6 localization in oocyst walls, sporozoites, and residual bodies (RBs) in ruptured oocysts (left), compared with COWP4, which is largely restricted to the suture (right). **(B)** Distribution patterns of COWP6 in sporozoites during excystation. COWP6 localizes to the pellicle in unexcysted sporozoites and progressively redistributes to the crystalloid body (CB) and pre-nuclear region (PreN) during excystation. Quantification of distribution patterns over time is shown. **(C)** Relative abundance of COWP6 and COWP4 in oocyst walls and sporozoites based on published proteomic data. **(D)** Detection of secreted COWP6 in gliding sporozoites, showing deposition along gliding trails. **(E)** ELISA detection of secreted COWP6 in post-excystation medium and in supernatants collected during parasite invasion and intracellular development. PanCp antibody detection is shown as a reference. **(F)** Binding of recombinant COWP6-His (0.2 μM) to fixed HCT-8 cells, compared to GST-His control. **(G)** Binding kinetics of COWP6-His to host cells determined by cell-based ELISA (Kd = 0.037 μM). Inset shows SDS-PAGE of purified recombinant protein.

During excystation, COWP6 exhibited dynamic redistribution within sporozoites (Fig. 6B). In unexcysted sporozoites, COWP6 was predominantly localized to the pellicle. As excystation progressed, increasing proportions of sporozoites showed additional localization of COWP6 to the crystalloid body (CB) and a pre-nuclear region. These changes in distribution over time are consistent with active trafficking or redistribution of COWP6 during excystation. The presence of COWP6 in sporozoites is consistent with published proteomic data, which show that COWP6 is detected at substantially higher abundance in sporozoites than in oocyst walls, whereas the suture-specific COWP4 is present at much lower levels (Fig. 6C) [19–21].

These observations prompted us to investigate whether COWP6 is secreted by sporozoites. In a gliding motility assay, COWP6 was detected as granular deposits along sporozoite trails, indicating secretion during motility (Fig. 6D). Secretion of COWP6 was further confirmed by ELISA, which detected COWP6 in post-excystation medium and in culture supernatants collected during early invasion (0–3 hpi) and subsequent intracellular development (Fig. 6E).

The presence of COWP6 in culture supernatants appears inconsistent with its lack of detectable signal in asexual stages by IFA (Fig. 3A). This discrepancy can be explained if COWP6 expressed during asexual stages is rapidly secreted, resulting in low intracellular accumulation. This interpretation is supported by transcript analysis (S6 Fig), which shows that COWP6 mRNA is present not only in oocysts and sporozoites but also at comparable levels during early intracellular stages (6–24 hpi) and later stages (48–72 hpi), whereas COWP4 transcripts are largely restricted to sporozoites and later developmental stages.

Given that COWP6 contains two GFR-CR domains with predicted adhesive properties, we tested whether it can bind host cells. Recombinant His-tagged COWP6 (COWP6-His) bound strongly to fixed HCT-8 cells in a cell-based ELISA assay, whereas no binding was observed for the GST-His control (Fig. 6F). Binding analysis revealed a high affinity interaction with an apparent dissociation constant (K_d_) of 37 nM (Fig. 6G).

Together, these results indicate that COWP6 is a multifunctional protein that serves not only as a structural component of the oocyst wall inner layer but also as a secreted adhesin capable of interacting with host cells.

### COWP6 directly interacts with COWP4

Given the adhesive properties of COWP6, its co-localization with COWP4 at the suture, and its relative resistance to proteolytic removal (Fig. 1F), we hypothesized that COWP6 may contribute to oocyst wall architecture by interacting with COWP4. To test this hypothesis, we performed bimolecular fluorescence complementation (BiFC) and far-western blot assays. In the BiFC assay, COWP6 and COWP4 were fused to the N- and C-terminal fragments of Venus fluorescent protein (COWP6–VN173 and COWP4–VC155). Strong fluorescence signals were observed in cells co-expressing these constructs, comparable to the positive control pair (bJun/bFos), whereas no signal was detected in negative controls, including empty vectors and mismatched pairs (Fig. 7A,B).

**Fig. 7.**
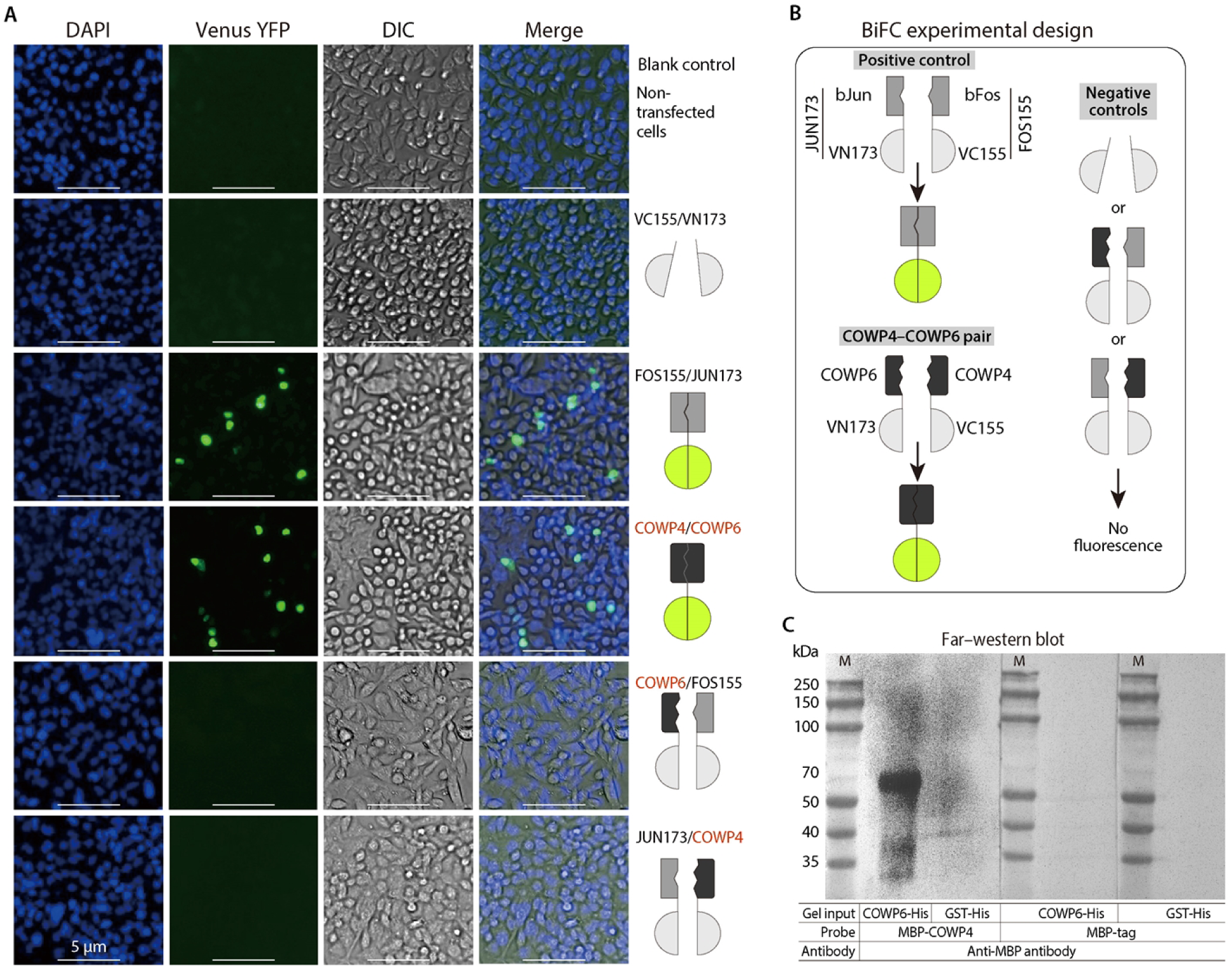
Direct interaction between COWP4 and COWP6. **(A)** Bimolecular fluorescence complementation (BiFC) assay demonstrating interaction between COWP4 and COWP6. Fluorescence signals are observed only in cells expressing the COWP4–COWP6 pair and the positive control (bJun/bFos), but not in negative controls. **(B)** Schematic of BiFC experimental design. **(C)** Far-western blot analysis showing binding of MBP-tagged COWP4 to immobilized COWP6-His, but not to GST-His. MBP alone does not bind either protein.

In far-western blot assays, MBP-tagged COWP4 specifically bound to immobilized COWP6-His but not to GST-His (Fig. 7C). Conversely, MBP alone did not bind to either protein, confirming the specificity of the interaction.

Collectively, these results demonstrate that COWP4 and COWP6 can directly and specifically interact both in vitro and in a cellular context, supporting a model in which COWP6 contributes to suture organization through interaction with COWP4.

## DISCUSSION

The *Cryptosporidium* oocyst wall protects the enclosed sporozoites and is essential for parasite survival in the environment, making it a key target for interrupting transmission. Although several Cryptosporidium oocyst wall proteins (COWPs) and putative oocyst wall proteins (POWPs) were identified over two decades ago, only COWP1 and COWP8 were experimentally localized to the inner and central wall layers, respectively [9,12,13]. Progress in this area remained limited until a recent study employing CRISPR/Cas9-mediated tagging of COWP2–COWP9 revealed their subcellular distributions within the oocyst wall (Bacchetti et al., 2025) [14]. That study showed that COWP2–COWP4 localize to the oocyst suture, whereas COWP5–COWP9 are distributed in the inner wall layer, and further demonstrated that COWP8 is dispensable for oocyst formation and transmission. While these findings significantly advanced our understanding of oocyst wall composition, key questions remain regarding the functional roles of individual COWPs, their interactions, and the molecular mechanisms governing suture structure and opening.

In the present study, we demonstrate that the suture-specific protein COWP4 plays an essential role in parasite transmission. Disruption of the *COWP4* gene in *C. parvum* abolishes excystation and prevents infection of new hosts, establishing COWP4 as a critical determinant of infectivity. Notably, COWP4 deletion does not impair intracellular development, as parasites are still able to complete the life cycle and produce morphologically normal oocysts in IFN-γ-KO mice. However, these oocysts represent a dead-end stage due to their inability to excyst and initiate subsequent infection. These findings provide direct functional evidence that the oocyst suture is a genetically defined gatekeeper for parasite transmission.

In addition to its role in excystation, COWP4 deletion markedly reduces oocyst viability during storage, suggesting that the suture may contribute not only to sporozoite release but also to maintaining oocyst integrity under environmental conditions. While the underlying mechanisms remain unclear, the loss of COWP4 may compromise the structural stability of the suture region, rendering oocysts more susceptible to environmental stress. Further studies will be required to define how suture architecture contributes to long-term oocyst survival.

Beyond its structural role, COWP6 is also present in sporozoites and is secreted during excystation, motility, and host cell invasion. The protein exhibits high-affinity binding to host cells in the low nanomolar range, supporting a role as a secreted adhesin. Although its precise biological function remains to be determined, COWP6 may contribute to parasite attachment, invasion, or modulation of host cell responses. The multifunctional nature of COWP6 is further supported by the inability to generate COWP6-null parasites, suggesting that it may be essential for parasite viability. Future studies employing conditional knockdown or domain-specific mutagenesis will be important to dissect its roles during different stages of the parasite life cycle.

Together, these findings support a model in which the oocyst wall is composed of structurally and functionally specialized proteins with non-redundant roles. In this framework, COWP4 functions as a suture-specific gatekeeper that is essential for excystation and transmission, whereas COWP6 acts as a multifunctional protein linking oocyst wall architecture to parasite–host interactions. More broadly, this work provides new insights into the molecular organization of the oocyst wall and highlights the suture as a critical and genetically defined structure in *Cryptosporidium* biology.

From a translational perspective, both COWP4 and COWP6 represent potential targets for interrupting parasite transmission. Inhibiting COWP4 function could prevent excystation and block infection at an early stage, while targeting COWP6 may interfere with both oocyst wall integrity and host cell interactions. Given the environmental resilience of *Cryptosporidium* oocysts and the limited efficacy of current control measures, disruption of oocyst wall structure and function represents a promising strategy for controlling transmission. Further investigation of oocyst wall proteins and their interactions may therefore provide new avenues for therapeutic or environmental intervention.

## Conclusions

This study provides a functional dissection of the *Cryptosporidium* oocyst suture and demonstrates that the suture-specific protein COWP4 is essential for excystation, infectivity, and oocyst survival. In contrast, COWP6 is a multifunctional protein that contributes to oocyst wall architecture, is secreted during parasite motility and invasion, and directly interacts with host cells and COWP4. These findings reveal that the oocyst wall is composed of structurally and functionally specialized proteins with non-redundant roles, and establish the suture as a genetically defined gatekeeper for parasite transmission. Targeting oocyst wall components, particularly those involved in suture function, may represent a promising strategy for interrupting the transmission cycle of *Cryptosporidium*.

## Materials and methods

### Ethics statement

All animal procedures complied with the *Guide for the Care and Use of Laboratory Animals* of the Ministry of Health of China. Animal use protocols were approved by the Animal Welfare and Research Ethics Committee of the Institute of Zoonosis, Jilin University (AUP #2020-IZ-20, #2024-IZ-00031, and #2024-IZ-00033).

### Parasites and cell lines

*Cryptosporidium parvum* (gp60 subtype IIaA17G2R1) oocysts were harvested from the feces of experimentally infected calves and purified by sucrose gradient centrifugation as previously described [22,23]. Purified oocysts were resuspended in phosphate-buffered saline (PBS) containing antibiotics (100 U/mL penicillin and 100 μg/mL streptomycin) and stored at 4 °C.

For excystation and in vitro culture, oocysts were surface-disinfected with 0.2% sodium hypochlorite (5 min on ice), followed by at least five washes with PBS. Excystation was performed by incubating oocysts in RPMI-1640 medium containing 0.75% sodium taurocholate at 37 °C for 30–45 min. Only oocysts with >80% excystation efficiency were used. Excysted sporozoites were immediately used for in vitro infection, immunofluorescence assay (IFA), transmission electron microscopy (TEM), or protein localization analyses.

For in vitro culture, HCT-8 cells (ATCC #CCL-244) were maintained in RPMI-1640 supplemented with 10% fetal bovine serum (FBS), penicillin (100 U/mL), and streptomycin (100 μg/mL) at 37 °C with 5% CO₂. For IFA, cells were grown on poly-L-lysine-coated glass coverslips in 48-well plates to appropriate confluency depending on the experiment [24]. Bleach-treated oocysts or freshly excysted sporozoites were added at parasite-to-cell ratios of approximately 2:1 (oocysts) or 4:1 (sporozoites). Samples were collected at specified time points for IFA, TEM, or protein localization analyses [24,25].

### Antibody preparation

Peptides unique to *C. parvum* COWP4 (cgd8_3350; EGGIEPDSDNMC, amino acids 440–451) and COWP6 (cgd4_3090; CESKIHEKHHGKNTRT, amino acids 518–533) were synthesized (China Peptides, Shanghai, China) and conjugated to keyhole limpet hemocyanin (KLH) [26]. Polyclonal antibodies were generated in New Zealand white rabbits using standard immunization protocols.

Primary immunization was performed with 300 μg of KLH-conjugated peptide emulsified in complete Freund’s adjuvant via multiple subcutaneous injections. Booster immunizations were performed with 150 μg peptide in incomplete Freund’s adjuvant on days 14, 28, and 42. Serum samples were collected prior to immunization (pre-immune) and 7 days after the final boost.

Antibodies were affinity-purified using peptide-coated nitrocellulose membranes as previously described [27,28]. Briefly, 100 μg peptide was applied to nitrocellulose membranes and air-dried. Membranes were blocked for 1 h, washed with TBST (0.15 M NaCl, 0.01 M Tris-HCl, 0.05% Tween-20, pH 7.4), and incubated with diluted antiserum (0.2 mL serum in 2 mL 0.1% BSA/TBST) for 2 h. After washing, bound antibodies were eluted with glycine buffer (0.2 M glycine, 0.15 M NaCl, 0.05% Tween-20, pH 2.7) and immediately neutralized with 1.0 M Tris (pH 10). Purified antibodies were used immediately or stored at 4 °C (short-term) or −20 °C (long-term).

### Western blot analysis

Oocysts (1–2 × 10^7^ per lane) were lysed in 20 μL RIPA buffer containing protease inhibitors using three freeze–thaw cycles between liquid nitrogen and ice, followed by centrifugation to remove insoluble material. Supernatants were separated by SDS-PAGE and transferred to PVDF membranes.

Membranes were blocked with 5% BSA in TBST and incubated with affinity-purified COWP4 or COWP6 antibodies (1:10 dilution in 5% skim milk/TBST), followed by HRP-conjugated goat anti-rabbit IgG (1:5000; Invitrogen). Signals were visualized using enhanced chemiluminescence (ECL) (Beyotime) and imaged using a ChemStudio-815 system (Analytik Jena). Unless otherwise specified, incubations were performed for 1 h at room temperature with three 5-min washes between steps.

### Indirect immunofluorescence assay (IFA)

IFA was used to localize COWP4 and COWP6 in oocysts, sporozoites, and intracellular stages. To resolve localization within oocyst wall layers, the following preparations were used: 1) Intact oocysts with outer veil (OV) preserved; 2) OV-depleted oocysts treated with sodium hypochlorite; 3) OV-depleted oocysts subjected to controlled digestion with pronase E (0.7 U/mL, room temperature for 24–48 h); 4) Excysted oocyst wall “ghosts”; and 5) Ghost walls subjected to additional pronase digestion.

Samples were fixed in 4% paraformaldehyde (30 min), washed, and applied to poly-L-lysine-coated slides. After drying, samples were permeabilized with 0.2% Triton X-100 (5 min), blocked with 3% BSA (1 h), and incubated with primary antibodies (1:10 dilution) followed by Alexa Fluor 488-conjugated secondary antibodies. Samples were counterstained with DAPI and mounted for imaging using an Olympus BX53 microscope.

For intracellular stages, infected HCT-8 cells were fixed and processed similarly.

For dual labeling, COWP4 antibodies were pre-labeled with Alexa Fluor 488. Samples were first stained with anti-COWP6 antibody and Alexa Fluor 594 secondary antibody, followed by Alexa Fluor 488-conjugated COWP4 antibody.

### Recombinant protein expression and purification

Signal-peptide-truncated *C. parvum* COWP4 (aa 23 to 887) was expressed as maltose-binding protein (MBP)-tagged fusion protein (MBP-CpCOWP4; or rCpCOWP4). Signal-peptide-truncated *C. parvum* COWP6 (aa 23 to 538) was expressed as His(6×)-tagged fusion protein (His-CpCOWP6; or rCpCOWP6). Gene fragments were amplified from *C. parvum* genomic DNA by PCR Phanta Max Super-Fidelity DNA Polymerase (Vazyme, Nanjing, China), with the following primers: COWP4-SalI-F (5′-acgc<u>GTCGAC</u>AAACCTGGAATTTCGGCATCAC-3′) and COWP4-HindIII-R (5′-ccc<u>AAGCTT</u>TTATAAATTCAGATTTGTTCTAATGTTTG-3′) for CpCOWP4, and COWP6-BamHI-F (5′-cgc<u>GGATCC</u>CAGCACGGACAACATTTG-3′) and COWP6-XhoI-R (5′-cc<u>CTCGAG</u>TTAATAGTGGCTATTTGCAGTCCTA-3′) for CpCOWP6. Lowercase letters indicate protective bases, and underlined sequences denote restriction sites.

COWP4 and COWP6 amplicons were cloned into pMAL-c5X (New England Biolabs) and pET-28a (Invitrogen), respectively. MBP alone and His-GST proteins were also expressed for use as negative controls. After sequence verification, constructs were transformed into *Escherichia coli* Rosetta(DE3) strain competent cells. In a typical expression experiment, 1000 mL of bacterial suspension was grown in LB broth at 37 °C until OD_600_ reached to ∼0.6, followed by the addition of isopropyl-β-D-thiogalactopyranoside (IPTG; 0.1 mM) to induce the expression at 16 °C for 10 h. Bacteria were harvested by centrifugation, and resuspended in 20 mL of PBS containing 1% Triton X-100 and phenylmethylsulfonyl fluoride (PMSF), and lysed by sonication on ice. Lysates were centrifuged (12,000 × g for 30 min) to collect pellets. The recombinant proteins were affinity-purified with amylose-resin chromatography for rCpCOWP4 or immobilized nickel-resin chromatography (rCpCOWP6) following standard protocols. Eluted proteins were pooled and dialyzed in PBS. The quality and quantity were assessed by SDS-PAGE and Bradford assay, respectively.

### Interaction between COWP6 and COWP4 by bimolecular fluorescence complementation (BiFC) and far-western blot (FWB) assays

BiFC was performed using split Venus fluorescent protein to evaluate protein interaction in living cells [29]. Four plasmids were acquired from Addgene (Watertown, MA, USA): pBiFC-VN173 (Addgene plasmid #22010) and pBiFC-VC155 (#22011) were used to construct COWP6-VN173 and COWP4-VC155 fusions; while pBiFC-bJunVN173 (#22012) with bJun-VN173 fusion and pBiFC-bFosVC155 (#22013) with bFos-VC155 fusion were used as positive control pair.

COWP6 (aa 23–538) and COWP4 (aa 23–887) were fused into VN173 and VC155, respectively. Gene fragments were amplified from *C. parvum* genomic DNA using the following primers: COWP6-EcoRI-F (5’-cg<u>GAATTC</u>CAGCACGGACAACATTTG-3’) and COWP6-KpnI-R (5’-gg<u>GGTACC</u>TTAATAGTGGCTATTTGCAGTCCTA-3’). The cloning of COWP4 (aa 23–887) into pBiFC-VC155 used primers COWP4-SalI-F (5’-acgc<u>GTCGAC</u>AAACCTGGAATTTCGGCATCAC-3’) and COWP4-XhoI-R (5’-cc<u>CTCGAG</u>TTATAAATTCAGATTTGTTCTAATGTTTG-3’). Lowercase letters indicate protective bases and underlined sequences denote restriction sites. The resulting constructs were termed COWP4-VC155 and COWP6-VN173.

Specified pairs of constructs were transfected into HCT-8 cells cultured on glass coverslips in 24-well plates with ∼70% confluence. Transfection was mediated by incubating cells with a premixture of plasmids (250 ng each) and polyethylenimine (PEI) (1 µg) in 200 μL of serum-free RPMI-1640 medium at 37 °C for 6 h. Transfection medium was replaced with FBS-containing medium, and cells were cultured for additional 24–36 h. Cells were fixed in 4% paraformaldehyde for 15 min at room temperature, washed three times with PBS, counterstained with DAPI (1 µg/mL, 10 min), mounted onto slides with antifade mounting medium, and examined under an Olympus BX53 microscope. For each condition, at least five random fields were collected, and the percentage of cells displaying specific BiFC fluorescence was quantified relative to the controls.

Far-western blotting was performed using immobilized His-COWP6 probed with MBP-COWP4, followed by detection with anti-MBP antibodies. Recombinant His-COWP6 (1–2 µg per lane) was resolved by SDS-PAGE, transferred to PVDF membrane, and blocked in 5% BSA/TBST. The membranes were probed with 50 μg of recombinant MBP-COWP4, or MBP-tag as a negative control, in 4 mL TBST for 1 h. After three washes with TBST, the membranes were incubated with anti-MBP monoclonal antibody (Abclonal, Wuhan, China) at 4 °C overnight, followed by incubation with horse radish peroxidase (HRP)-conjugated secondary antibody and visualization ECL reagents as described for western blot analysis above.

### Binding kinetics of COWP6 to host cells

For IFA, HCT-8 cells grown to confluence on coverslips (48-well plates) were fixed in cold 4% PFA in PBS for 30 min, washed with PBS, and blocked with 5% skim milk in PBS for 1 h at room temperature. Cells were incubated with COWP6-His (0.2 µM) or GST-His control (0.2 µM) in high-salt binding buffer (PBS containing 1.0 mM CaCl_2_, 1.0 mM MnCl_2_, and 500 mM NaCl) overnight at 4 °C. After three 5-min washes with PBS, cells were re-fixed in cold methanol for 10 min, blocked for 1 h, and incubated with a mouse anti-His antibody (Epizyme Biotech, China, 1:5000 dilution) and donkey anti-mouse IgG antibody conjugated with Alexa Fluor 488 (Invitrogen; 1:2000 dilution). Nuclei were counterstained with DAPI (1 μg/mL). Samples were examined under Olympus BX53 microscope.

For cell-based ELISA, HCT-8 cells cultured in 96-well plates to confluence were fixed with 1% glutaraldehyde in PBS for 30 min, washed three times with PBS, and blocked with 5% skim milk in PBS for 1 h at 37 °C. GST-COWP6 or GST-His in 500 mM NaCl/PBS at serial concentrations (0–2 µM) overnight at 4 °C. After three washes in PBS, cells were incubated with a mouse anti-His antibody and an alkaline phosphatase-conjugated goat anti-mouse IgG (H+L) (ImmunoWay Biotechnology; 1:10,000 dilution). Color development was performed using p-nitrophenyl phosphate (pNPP, Sigma-Aldrich) following the manufacturer’s protocol, and the optical density at 405 nm was measured using a multifunctional microplate reader (BioTek, Winooski, VT, USA).

### General procedure in mouse infection experiments

Interferon-γ-knockout (IFN-γ-KO, or GKO) mice with C57BL/6J background was used for infection with *C. parvum* to prepare in vivo samples for IFA detection of COWP4 and COWP6 in various developmental stages. IFN-γ-KO mice were also used for in vivo selection and evaluation of COWP4-KO parasite (see below).

For in vivo samples, five 6-week-old female mice were orally inoculated with wild-type (WT) *C. parvum* oocysts (5×10^4^ oocysts/mouse). Mice were sacrificed on specified days. Ileal samples were collected, gently rinsed with PBS to remove lumen contents, cut into 0.5-1cm segments, fixed with 10% neutral buffered formalin for 24 h, and washed three times with PBS (5-min each). Tissue specimens were paraffin-embedded, sectioned (4–5 µm), deparaffinized and dehydrated, following standard histological protocol. Sectioned tissues were treated with citrate buffer (pH 6.0) at 95°C for 10 min and then cooled to room temperature to inactivate endogenous peroxidase activity.

For IFA, permeabilization, blocking, labeling with primary (i.e., rabbit anti-COPW4/COWP6) and secondary (i.e., Alexa Fluor 488- or 594-conjugated goat anti-rabbit IgG) antibodies, counterstaining with DAPI, and microscopic examination used the same procedure as described above for IFA.

For hematoxylin and Eosin (H&E) staining and histological analysis, distal ileum segments were collected, rinsed with PBS, and fixed in 4% PFA at 4 °C for 24 h, followed by routine dehydration, paraffin-embedding, and sectioning (4–5 µm). Sections were deparaffinized in xylene, rehydrated through a graded ethanol series, stained with hematoxylin and eosin (H&E), dehydrated, mounted, and examined under a light microscope.

### Generation of *C. parvum* COWP4-KO

A CRISPR/Cas9 system was used to disrupt the COWP4 locus by inserting an mNeonGreen-Nluc-Neo expression cassette [30,31]. A single guide RNA (sgRNA; 5’-GTCGTACTGGTGATTTGTGC-3’) targeting a site near the N-terminus of the coding region was cloned into hSpCas9/U6 vector (see Fig. 6A for illustration). A donor repair template (donor DNA cassette) was generated by amplifying the mNeonGreen-Nluc-Neo cassette flanked by 60 bp homology arms corresponding to sequences upstream and downstream of the sgRNA-targeted site to drive homologous recombination. The donor DNA cassette for insertion contained fluorescent protein mNeonGreen for visualization, nanoluciferase (Nluc) for sensitive chemiluminescence detection, and neomycin-resistant marker (Neo) for in vivo selection with paromomycin.

The CRISPR/Cas9 plasmid (50 µg) and donor DNA (50 µg) were mixed with 1×10^8^ freshly excysted sporozoites in 75 μL of SF Cell Line 4D-Nucleofector X Kit buffer. Electroporation was conducted with Amaxa 4D Nucleofector system (program EH-100). Electroporated sporozoites were divided into four aliquots for oral inoculation into four IFN-γ-KO mice, which were pretreated with 0.8% sodium bicarbonate to neutralize stomach acid. At 18 h post-infection (hpi), paromomycin (16 g/L) was supplied in drinking water for in vivo selection. Fecal samples were collected from 3 dpa daily or every other day for up to 25 days for monitoring luciferase activity in fecal samples as described below.

Nanoluciferase activity in fecal samples was evaluated using NanoLuc Luciferase Assay System (Promega, Madison, WI, USA). Fecal pellets (50 mg) were soaked in 0.5 mL of lysis buffer, and subjected to five freeze-and-thaw cycles between liquid nitrogen and ice, followed by vigorous vortexing and centrifugation at 10,000 × g for 2 min. Supernatant (100 µL) was transferred to a 96-well plate, and mixed with 100 µL of diluted substrate-buffer mixture (1:50 dilution). Luminescent signals were measured immediately using a multimode microplate reader. Data were expressed as relative luciferase activity per mg feces, using feces from uninfected mice as a blank control.

Oocysts were purified from feces positive with luciferase activity by sucrose (specific gravity 1.33) and cesium chloride gradient centrifugations (specific gravity 1.15) as described [30,31]. COWP4 ablation was confirmed by PCR detection of the two junction sites of the integrated cassette at the COWP4 locus using genomic DNA isolated from COWP4-KO and wild-type oocysts and the following two primer pairs: 5’-AATTGTCAAACTGCTCTCGC-3’ and 5’-CAAGGTGCATGCAATTTGAC-3’ flanking the 5’-junction (1,145 bp) and (767 bp; 5’-CTGAAGCCGGTAGAGACTG-3’ and 5’-CTCCGGACACTCAATGTTTG-3’ flanking the 3’-junction (767 bp). PCR products were resolved on 1% agarose gels. The deletion of COWP4 was also evaluated by IFA and western blot of COWP4-KO oocysts in comparison with WT oocysts using routine protocols.

### Assessment of COWP4-KO oocysts for viability and infectivity

The in vitro excystation of COWP4-KO oocysts (vs. WT oocysts) was examined as described for the preparation of free sporozoites above. Its effect on the viability of oocysts stored over time at 4 °C was evaluated by one-step SYBR Green qRT-PCR targeting 18S rRNA transcript levels was performed (vs. WT oocysts) [5,32]. The viability of oocysts stored at 4 °C for 90 days was further evaluated by ATP assay using BacTiter-Lumi Microbial Cell Viability Assay Kit (Beyotime, Shanghai, China) as instructed by the manufacturer.

The in vivo infectivity of COWP4-KO was assessed in IFN-γ-KO mice in comparison to a thymidine kinase-knockout (ΔTK) strain of *C. parvum*, in which the TK locus was interrupted by the same mNeonGreen-Nluc-Neo cassette [31]. COWP4-KO and ΔTK oocysts (5×10^4^ per mouse) were inoculated. Fecal samples were collected for quantifying the oocyst shedding by nanoluciferase assay as described. Mice were sacrificed on 21 day post-infection (dpi), and small intestinal tissues were collected for histology, IFA, and electron microscopy.

### Transmission electron microscopy (TEM) and immunogold electron microscopy (IEM)

IEM was conducted for colloidal gold-labeling of COWP4 and COWP6 on ghost oocyst walls. Oocyst wall shells after excystation were applied onto nickel grids, followed by air-dry, three washes with PBS and blocking in PBS containing 5% skim milk and 0.01% Tween-20. Grids were incubated overnight at 4 °C with affinity-purified anti-COWP4 or anti-COWP6 antibodies (1:2 dilution), washed with PBS, and incubated with 10-nm colloidal gold-conjugated goat anti-rabbit IgG (Sigma-Aldrich) for 1 h at 37 °C. After three washes with PBS, grids were stained with 2% uranyl acetate and examined on a Hitachi H7650 transmission electron microscope. Gold particles were quantified in defined regions, such as suture edges and other areas. Images were captured with AMT XR40B CCD camera.

TEM was conducted to examine distal ileal tissues from GKO mice, uninfected or infected with WT or COWP4-KO oocysts. Ileal tissues on the epithelial side were cut into ∼0.5 mm^3^ pieces, and fixed for 2 h in PBS containing 2% PFA and 0.1% glutaraldehyde as described [33]. Samples were dehydrated through gradually increased ethanol (30%, 50%, 70%, 80%, 90%, and 100%; 1 h each), infiltrated with LR White resin (Sigma-Aldrich) at −20 °C for 48 h, and polymerized under UV light at −15 °C for 24 h. Ultrathin sections were prepared with a Leica EM UC6 ultramicrotome equipped with diamond knife, mounted on formvar-coated grids, and stained with 2% uranyl acetate. Grids were examined under a Hitachi H7650 transmission electron microscope, with images captured with AMT XR40B CCD camera.

## Supporting information

**S1 Fig.** Structural models of *Cryptosporidium parvum* oocyst wall proteins COWP4 and COWP6 as predicted by AlphaFold 3 (https://alphafoldserver.com).

**S2 Fig.** Immuno-gold labeling of COWP4 and COWP6 in post-excystation C. parvum oocyst walls, showing larger raw electron micrographs used in Figure 2A.

**S3 Fig.** Micrographs of F1-generation COWP4-KO oocysts obtained following in vivo selection. Oocysts purified from mouse feces were examined by differential interference contrast (DIC; upper panel) and fluorescence microscopy to detect mNeonGreen signals (lower panel), confirming efficient integration of the cassette.

**S4 Fig.** Quantitative measurements of major and minor axes of COWP4-KO and WT oocysts. COWP4-KO oocysts are slightly more elongated, with a higher shape index compared to WT. Statistical significance was assessed by Student’s t-test.

**S5 Fig.** Scanning electron micrographs of COWP4-KO and WT oocysts. No apparent differences in surface morphology were observed. Most oocysts were partially deformed during preparation; representative intact or minimally deformed oocysts are shown.

**S6 Fig.** Transcript levels of COWP4 and COWP6 during *C. parvum* development. Relative mRNA levels of COWP4 and COWP6 in oocysts, sporozoites, and intracellular stages (6–72 hpi) were determined by qRT-PCR and normalized to 18S rRNA. CpEF1α was included as a reference gene. COWP6 transcripts are detected across multiple developmental stages, whereas COWP4 transcripts are enriched in oocysts, sporozoites, and late developmental stages.

## Acknowledgments

This study was supported by funding from the National Key Research and Development Program of China (Grant No. 2023YFD1801000 to G.Z.) and the National Natural Science Foundation of China (Grant No. 31772731 to J.Y.). The funders played no role in the design of the study, data collection and analysis, the decision to publish, or the preparation of the manuscript.

## Author contributions

**Conceptualization:** Xiaodong Wu, Jigang Yin, Dongqiang Wang, Guan Zhu.

**Data curation:** Xiaodong Wu, Dongqiang Wang, Guan Zhu.

**Formal analysis:** Xiaodong Wu, Dongqiang Wang, Guan Zhu.

**Funding acquisition:** Jigang Yin, Guan Zhu.

**Investigation:** Xiaodong Wu, Dongqiang Wang, Wei Qi, Peng Jiang, Ying Zhang, Di Zhang.

**Methodology:** Xiaodong Wu, Jigang Yin, Dongqiang Wang, Guan Zhu.

**Supervision:** Dongqiang Wang, Guan Zhu.

**Validation:** Dongqiang Wang, Guan Zhu.

**Visualization:** Xiaodong Wu, Jigang Yin, Dongqiang Wang, Guan Zhu.

**Writing – original draft:** Xiaodong Wu, Jigang Yin, Dongqiang Wang.

**Writing – review & editing:** Dongqian Wang, Guan Zhu.

